# ProtFlap: Structure-based Deep Learning Prediction of Protein Flexibility from Atomic Energy Networks

**DOI:** 10.64898/2026.11.13.766391

**Authors:** Humaira Lubna, Yoonjoo Choi

## Abstract

Protein flexibility is a key determinant of molecular recognition, catalysis, and regulation, yet remains difficult to characterise at scale. Molecular dynamics (MD) simulations provide detailed descriptions of time-dependent dynamics but are computationally expensive. Although recent AI-based structure prediction methods have substantially improved access to high-quality static protein structures, they inherently lack information regarding the dynamic fluctuations essential for functional understanding. There is therefore a need for scalable, structure-based computational approaches that can infer intrinsic protein flexibility without explicit dynamical sampling. We present ProtFlap, a graph neural network framework for predicting relative residue-level flexibility directly from static protein structures. ProtFlap represents proteins as residue interaction graphs and integrates empirical atomic energy features extracted from a single energy-minimised snapshot with structural and sequence-derived descriptors. Central to our approach is the hypothesis that the local atomic energy network encoded in a frozen protein structure contains sufficient information to estimate structural flexibility. Trained using atomic mobility labels, quantified via normalised root-mean-square fluctuation (RMSF) derived from short MD simulations, ProtFlap achieves robust and accurate flexibility prediction across diverse protein domains. Feature attribution analysis reveals that van der Waals–related energy terms are the dominant contributors to the prediction.

**Key Points:**

- ProtFlap is a graph neural network that predicts intrinsic residue-level protein flexibility.
- The framework evaluates single, static structures, bypassing computationally expensive molecular dynamics simulations.
- Predictions are physically grounded by integrating structural descriptors with empirical atomic energy terms.
- Feature analysis reveals that local thermodynamic constraints, primarily Van der Waals forces, dominate flexibility prediction.

## 1 Introduction

Proteins are dynamic molecules whose time-dependent conformational transitions govern molecular recognition, catalysis, signalling, and regulation. Structural fluctuations enable ligand binding, enzymatic adaptation, and protein–protein interactions [13, 40]. In antibodies, dynamics within complementarity-determining regions (CDRs) strongly influences antigen recognition [4, 11]. Enzymes also require local conformational rearrangements for efficient substrate binding and catalysis [12]. Despite its biological importance, protein flexibility remains difficult to quantify and predict accurately [37]. Directly observing these continuous kinetic pathways through experimental techniques remains profoundly difficult. While experimental structure determination and recent AI-based prediction methods such as AlphaFold [19] and RoseTTAFold [3] provide high-quality static structural models, they represent single conformational snapshots that inherently lack information regarding time-dependent structural motion.

Historically, the primary experimentally derived metric used to gauge the structural movement has been the B-factor (or temperature factor), which measures atomic positional uncertainty within a crystal lattice. While B-factors provide valuable structural data, they are frequently influenced by crystallographic packing forces, averaging effects, and refinement artefacts. These external constraints can obscure the true intrinsic mobility of the protein in solution [6, 38].

Consequently, molecular dynamics (MD) simulations have become the standard computational framework for explicitly modelling these atomic motions over time [16]. By simulating time-dependent dynamics, MD provides a measurable, time-averaged record of mobility (physical spatial displacement). The mobility is typically quantified via the root-mean-square fluctuation (RMSF), which measures the average spatial deviation of each atom from its mean position [5, 42]. Because RMSF explicitly captures thermal motions independent of crystal lattice constraints, it serves as a robust, physically grounded metric for structural movement.

However, explicit MD simulation remains computationally costly. In the context of protein engineering, rapid and reliable estimation of this atomic mobility is critical. For example, modulating the conformational landscape of an enzyme active site is essential for catalytic adaptation [12], and mapping local structural movement is crucial for engineering stable binding interfaces in protein therapeutics.

Therefore, rather than attempting to explicitly simulate exhaustive dynamic pathways, it is highly advantageous to develop methods capable of inferring a protein’s intrinsic flexibility, the underlying thermodynamic potential for conformational heterogeneity, directly from static structural data. Several computational approaches have been developed to predict protein flexibility directly from sequence or structure [44]. Sequence-based methods, including MEDUSA [41], FlexC [46], and PROFbval [35], estimate residue flexibility from evolutionary and physicochemical features. Among structure-based approaches, simple descriptors based on local packing density have proven particularly effective in estimating residue-level fluctuations. In particular, weighted contact number (WCN) quantifies the packing environment of each residue through distance-weighted inter-residue contacts, providing a computationally inexpensive measure of local structural constraints. Previous studies have demonstrated strong relationships between protein packing density, WCN, and crystallographic atomic displacement parameters, while related models have also been evaluated against MD-derived fluctuation measures [14, 27, 47].

These methods are computationally efficient but do not explicitly incorporate structural context or residue interactions [45]. Structure-based approaches such as RMSF-Net [36], DEFMap [29], and normal mode analysis frameworks including ProDy [48] utilise three-dimensional structural information to improve flexibility estimation. More recent generative approaches such as BioEmu [25] model conformational ensembles directly from learned structural distributions. Although these methods advance flexibility prediction, they frequently rely on coarse-grained representations or purely geometric features that do not fully capture the local thermodynamic driving forces governing residue motion. Scalable prediction of residue-level dynamics from static structures, while preserving the underlying physical principles of atomic interactions, remains an unmet need.

Here, we present ProtFlap, a graph neural network framework for predicting residue-level relative flexibility directly from static protein structures. ProtFlap represents proteins as residue interaction graphs and integrates sequence-derived and structural descriptors with atomic energy features — specifically, empirical force-field terms such as Van der Waals and electrostatic energies extracted from a single energy-minimised structural snapshot. The model predicts continuous normalised RMSF values derived from MD simulations, with binary classification applied as a secondary threshold-based interpretation.

Our approach is based on the hypothesis that the local energetic environment encoded within an effectively frozen, energy-minimised protein structure contains sufficient information to estimate relative residue mobility. ProtFlap therefore links local atomic energy distributions with residue-level flexibility without explicit dynamical sampling. Rather than predicting long-timescale conformational transitions, ProtFlap estimates relative local flexibility patterns within folded protein domains. This formulation provides a physically grounded, interpretable, and scalable method for flexibility prediction across large structural datasets.

## 2 Methods

### 2.1 Dataset Selection

The dataset for this study was obtained from the CATH database (v4.3), curated up to 2022 [23]. The initial dataset contained 29,687 protein domains. Redundancy was reduced by excluding domains sharing more than 60% sequence overlap or 40% sequence identity with other domains. Only domains derived from X-ray crystal structures with a resolution of better than 2.7Å were retained. Domains containing two or more consecutive missing residues were excluded, and a minimum domain length of 50 residues was required. After applying these filters, the final dataset comprised 7,878 domains.

For external validation, we prepared an independent test dataset from ATLAS [42]. We excluded structures containing fewer than 50 residues and PDB entries overlapping with our CATH-derived dataset, leaving 785 PDB structures. These structures were divided into domains according to the ECOD classification [7]. After the filtration process, 1,096 domains were retained for external evaluation. Among the selected structures, 28 contained 65 experimentally annotated disorder-associated regions, which were used for a separate analysis of prediction performance in these regions.

For comparison with NMR structural ensembles, 239 protein structures containing at least 30 deposited models and 100 residues were selected from the RCSB PDB. NMR-based RMSF was calculated from all deposited models in each ensemble. For MD-based RMSF, the first model of each ensemble was used as the initial structure for a 1-ns GROMACS simulation with the AMBER force field.

### 2.2 Model Training and Evaluation

The dataset of 7,878 protein domains was divided into training, validation, and independent test sets. A total of 10% of the domains (788) was reserved as an independent test set and excluded from all stages of model development and hyperparameter optimisation. The remaining 90% (7,090) of the dataset was used for model training and validation. To evaluate the robustness of the framework to input structure derivation, corresponding predicted structural models for all 788 domains in the independent test set were additionally generated using ColabFold [30].

To assess robustness across data partitions, the training-validation subset was further divided into ten independent random splits using a 90:10 ratio. Model performance was evaluated across all validation splits, and the final model was selected based on the lowest validation loss.

ProtFlap was trained to predict continuous normalised RMSF values using a regression objective. Smooth L1 loss was used during optimisation. Model optimisation and hyperparameter tuning were performed using Optuna [2]. The hyperparameter search included hidden dimensions, learning rate, dropout probability, activation function, residual connections, normalisation schemes, and alternative graph neural network architectures including GCN, GIN, and GraphSAGE (Supplementary Table S1).All model were trained using the Adam optimiser with a batch size of 16. Training was performed for up to 100 epochs with early stopping applied based on validation loss, using a patience of 20 epochs to prevent overfitting. A ReduceLROnPlateau learning-rate scheduler was also used to adaptively reduce the learning rate when validation performance plateaued.

Regression performance was evaluated using root-mean-square error (RMSE) and Pearson correlation coefficient. For secondary binary interpretation, predicted normalised RMSF values were thresholded at zero to classify residues as relatively flexible or relatively rigid.

### 2.3 Atomic Energy Calculation and Molecular Dynamics

For each protein structure, atomic energy calculations were performed using Tinker (v8.9.2) [34] following energy minimisation to remove steric clashes and optimise local geometry using the AMBER99SB force field [17] with the implicit solvent model (GB/HPMF) [26]. Bonded and non-bonded energy components were computed and averaged at the residue level to generate per-residue atomic energy features (Supplementary Table S2).

MD simulations were performed using OpenMM (v7.0) [9] with the same force field (AMBER99SB) and the explicit solvent model (TIP3P water) [17, 18]. Protein structures were prepared using PDBFixer [9], solvated, and energy-minimised prior to simulation. Simulations were carried out at 300 K and 1 bar using a Langevin thermostat and Monte Carlo barostat under periodic boundary conditions. Each simulation was run for 1 ns with a 2 fs timestep and repeated three times.

Residue-level flexibility was quantified using normalised RMSF values. Raw values were normalised within each protein using a Z-score transformation [36] (Fig.2A).

### 2.4 Graph Representation and Feature Encoding

We developed a residue-level Graph Convolutional Network (GCN) combined with a multilayer perceptron (MLP) (Fig. 1). Protein structures were converted into graphs in which residues served as nodes and spatial contacts defined the edges. Protein graphs were generated using the NetworkX library, together with BioPython for processing protein structural information [8, 15]. For model development, we used PyTorch as our deep learning framework and incorporated Scikit-learn for additional analytical tasks [32, 33]. Given the variability in protein lengths, we used the collate_fn function from PyTorch Geometric to efficiently batch graphs of differing sizes. This approach preserved graph topology during batch processing without requiring padding.

**Figure 1:**
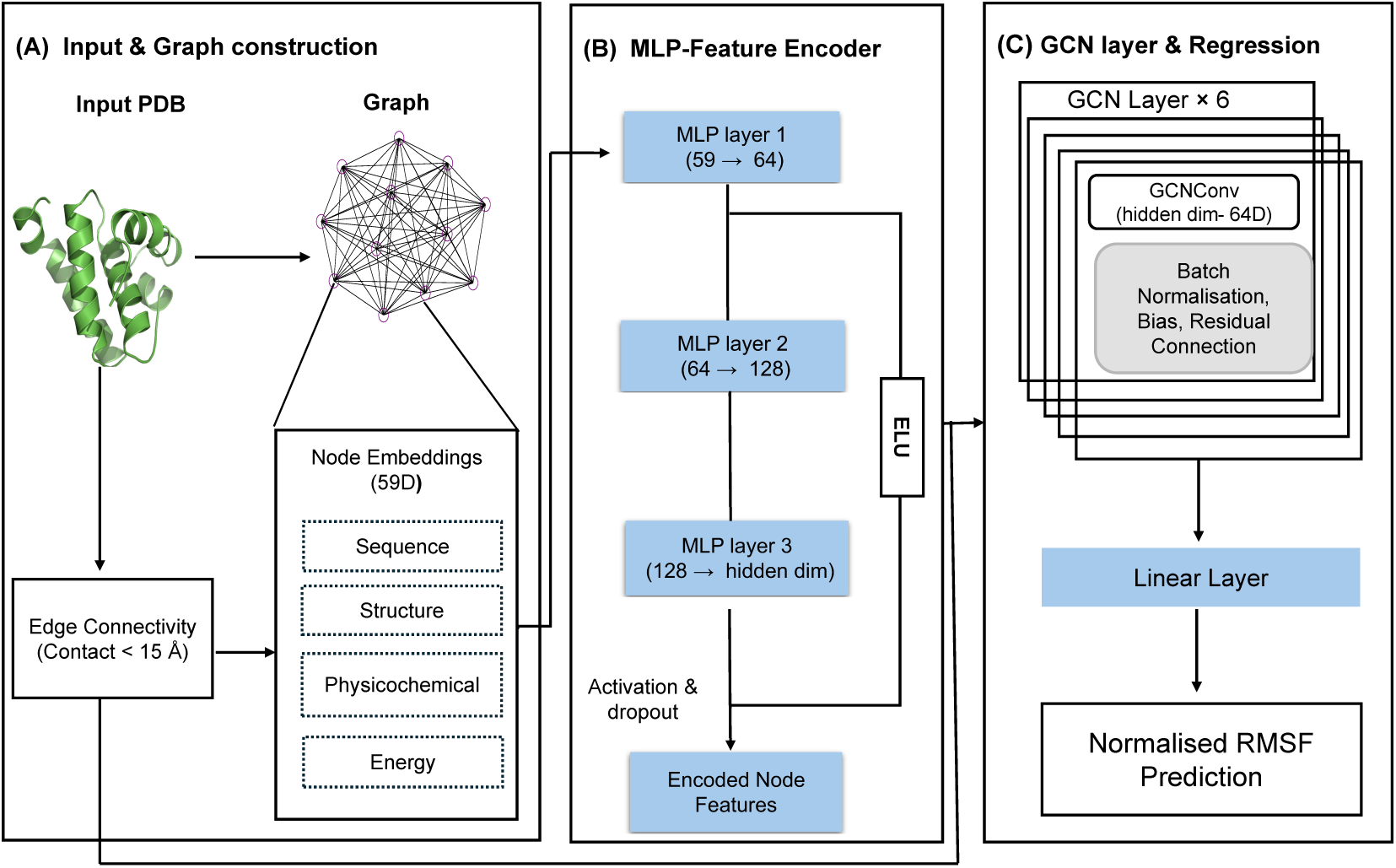
Overview of the ProtFlap framework. (A) **Input and graph construction.** Protein structures provided as PDB files are converted into residue interaction graphs. Each node represents an amino acid residue, and edges connect residues whose C*α* atoms lie within a 15 Å distance threshold. Edge weights are defined from inter-residue distances. Each node is annotated with a 59-dimensional feature vector comprising sequence-derived features, structural descriptors, physicochemical properties, and residue-level atomic energy terms derived from energy minimisation. (B) **MLP feature encoder.** Raw node features are processed through a three-layer multilayer perceptron (MLP) with ELU activation to generate latent node embeddings for graph message passing. (C) **GCN layers and regression.** Encoded node embeddings are processed through stacked edge-weighted graph convolutional network (GCN) layers with batch normalisation and residual connections. A final linear regression head outputs continuous normalised RMSF predictions for each residue.

The input node feature vector has 59 dimensions (Table 1), including 28 distinct atomic energy terms (Supplementary Table S2). The model consisted of an MLP feature encoder followed by stacked GCN layers and a final linear regression layer predicting normalised RMSF values. To optimise performance, we applied Optuna for hyperparameter tuning [2]. Edge weights derived from inter-residue distances were incorporated during message passing to enable weighted aggregation of neighbourhood information. The hyperparameter search space included hidden dimensions, activation functions, dropout probabilities, learning rates, optimiser settings, residual connections, normalisation schemes, and alternative GNN architectures including GCN, GIN, and GraphSAGE (Supplementary Table S1). The final model configuration was selected based on validation performance. Predicted normalised RMSF values were additionally thresholded to derive secondary binary flexibility labels.

**Table 1:**
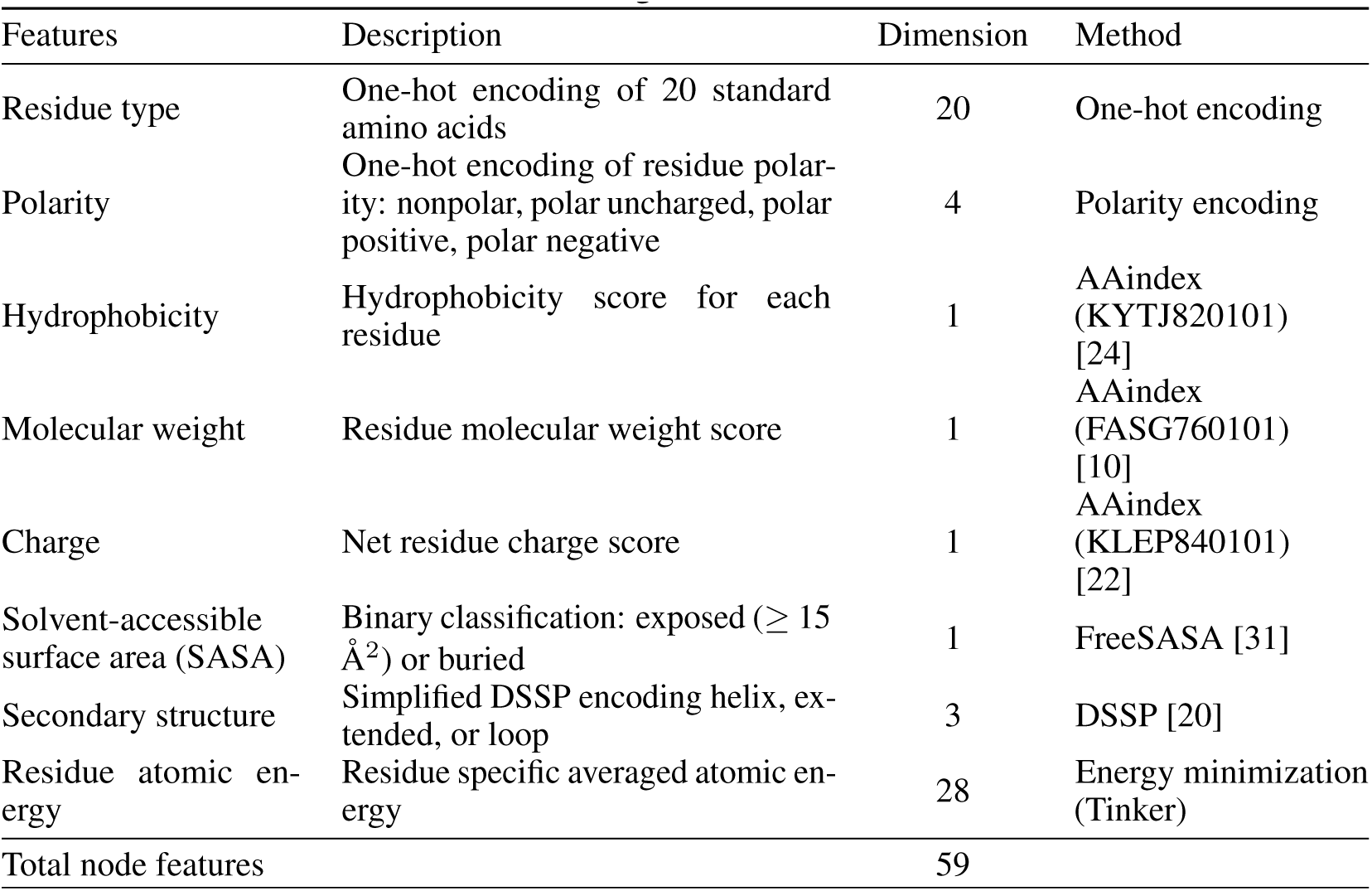
Feature categories and their dimensions.

Each protein structure was represented as a residue interaction graph in which nodes corresponded to amino acid residues and edges connected residues whose C*α* atoms were within 15 Å [43]. Node features included amino acid identity, physicochemical properties [10, 21, 22, 24], solvent accessibility [31], secondary structure assignments [20], and residue-averaged atomic energy terms.

### 2.5 Benchmark Methods and Comparative Evaluation

ProtFlap was benchmarked against representative structure-based and dynamics-based methods for protein flexibility estimation, including RMSF-Net [36], ProDy [48], BioEmu [25], and inverse weighted contact number (iWCN) [14, 27, 47]. Because these methods produce flexibility estimates using different physical representations and numerical scales, all outputs were converted into residue-level normalised values using per-protein Z-score normalisation prior to comparison.

For ProDy, residue fluctuation profiles were generated using the Gaussian Network Model (GNM) implemented in the ProDy framework. By default, the lowest 20 non-trivial normal modes were selected, as these low-frequency modes capture the dominant collective motions of proteins, whereas higher-frequency modes primarily represent local fluctuations. Mean square fluctuation (MSF) values were calculated using the calcSqFlucts function and subsequently converted to RMSF-like values. Because GNM-derived fluctuations are expressed in arbitrary relative units rather than absolute physical units, the resulting values were further normalised before comparison with MD-derived RMSF profiles.

For BioEmu, conformational ensembles were generated from each protein sequences, and residue-level RMSF values were calculated directly from the generated trajectories. The resulting RMSF profiles were subsequently normalised using the same procedure applied to ProtFlap predictions and MD-derived RMSF values.

RMSF-Net predictions were obtained from the published implementation. Because RMSF-Net outputs residue-level normalised flexibility values directly, no additional RMSF conversion was required.

For iWCN, weighted contact numbers were calculated from the C*α*–C*α* distances between each residue and all other residues in the protein structure, following the conventional inverse-square distance weighting:

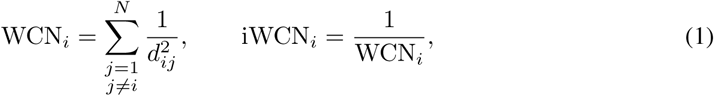

where *d_ij_* is the C*α*–C*α* distance between residues *i* and *j*, and *N* is the total number of residues in the protein. Higher iWCN values indicate lower local packing density and potentially greater residue flexibility. The resulting iWCN values were normalised using per-protein Z-score transformation before comparison with MD-derived RMSF profiles.

## 3 Results

### 3.1 Overview of the ProtFlap Framework

ProtFlap estimates residue-level flexibility directly from static protein structures. ProtFlap is a graph neural network framework designed to predict residue-level protein flexibility directly from static protein structures (Fig. 1). Protein structures are represented as residue interaction graphs in which nodes correspond to amino acid residues and edges connect residues with C*α*–C*α* distances below 15 Å. This cutoff was selected after evaluating five distance thresholds (5, 10, 15, 20 and 30 Å), with 15 Å giving the highest mean Pearson correlation (*r* = 0.65), compared with *r* = 0.62 at the original 30 Å cutoff (Supplementary Fig. S1).

The model combines a multilayer perceptron (MLP) feature encoder with stacked graph convolutional network (GCN) layers to propagate information across local structural environments. By integrating energetic and structural information within the graph representation, ProtFlap predicts continuous normalised RMSF values for individual residues.

### 3.2 Normalised RMSF Patterns Are Stable Across Simulation Conditions

To assess whether short MD simulations provide reliable relative flexibility labels, normalised RMSF profiles derived from the CATH dataset (1 ns) and the ATLAS dataset (1 and 100 ns) were first compared [42]. The probability density distributions (Fig. 2A) showed close agreement, indicating similar overall distributions of relative residue mobility.

**Figure 2:**
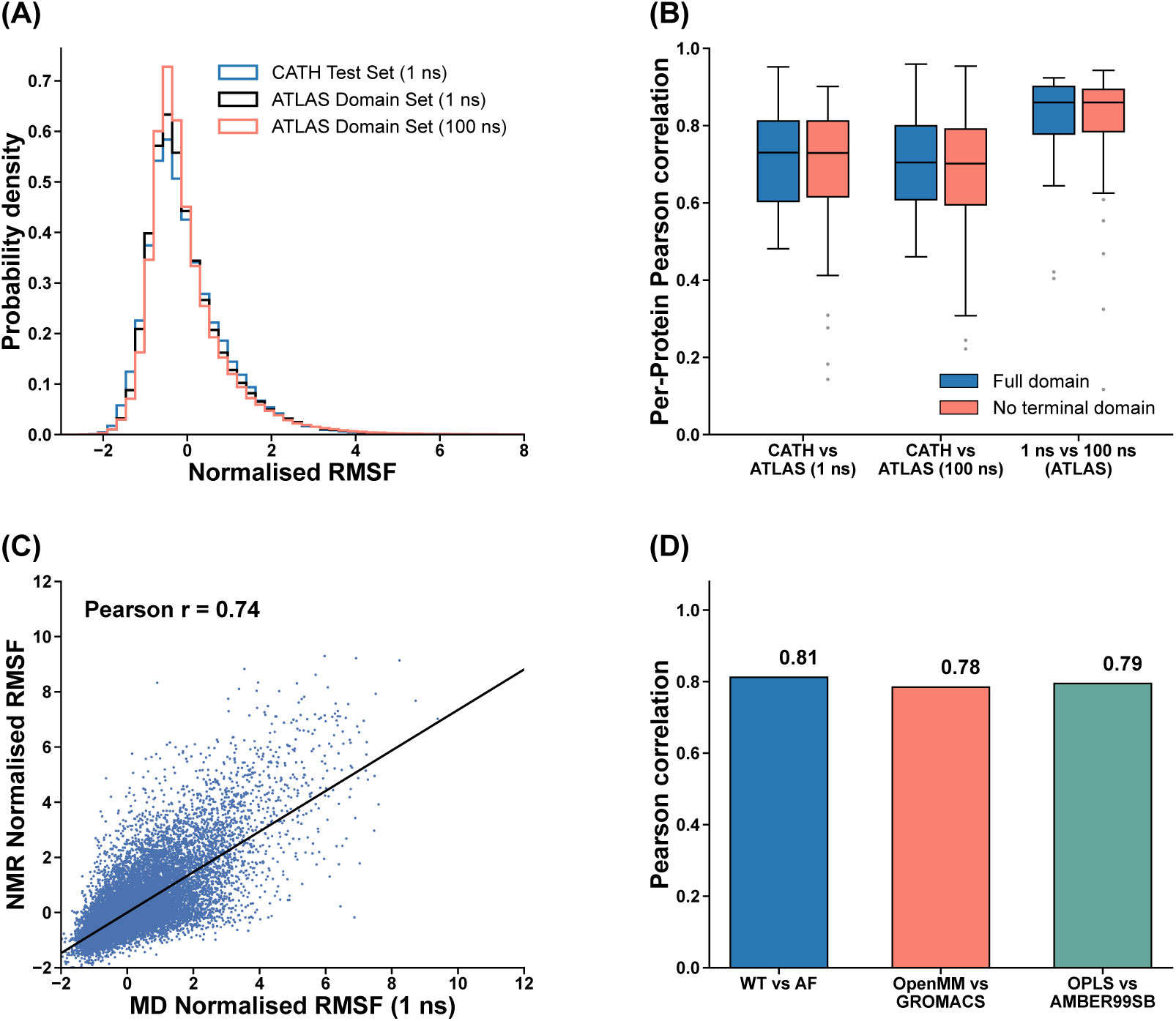
Consistency of normalised RMSF across simulation lengths, structural sources, and MD protocols. (A) Probability density distributions of normalised RMSF from the CATH test set (1 ns) and ATLAS domain set (1 and 100 ns). (B) Distributions of per-protein Pearson correlations between CATH 1 ns and ATLAS 1 ns, CATH 1 ns and ATLAS 100 ns, and ATLAS 1 ns and ATLAS 100 ns. Correlations were calculated using all residues (full domain) and after excluding the first and last 10 residues. The corresponding correlations were 0.71/0.70, 0.70/0.69, and 0.84/0.83, respectively (full domain/without terminal residues). (C) Comparison of normalised RMSF derived from NMR structural ensembles and 1-ns MD simulations (Pearson *r* = 0.74). (D) Pearson correlations between normalised RMSF profiles obtained using different structural sources and MD protocols: experimental crystallographic structures (WT) versus AlphaFold2-derived models (AF) (*r* = 0.81), OpenMM versus GROMACS using identical AMBER parameters (*r* = 0.78), and OPLS versus AMBER99SB force fields within GROMACS (*r* = 0.79).

To determine whether this agreement was consistent across individual proteins, we calculated per-protein Pearson correlations between normalised RMSF profiles from the CATH and ATLAS domain datasets. Comparisons were performed between CATH 1 ns and ATLAS 1 ns (Pearson *r* = 0.71), CATH 1 ns and ATLAS 100 ns (Pearson *r* = 0.70), and ATLAS 1 ns and ATLAS 100 ns (Pearson *r* = 0.84) using matched protein structures (Fig. 2B). To assess whether the correlations were dominated by highly flexible terminal regions, the analysis was repeated after excluding the first and last 10 residues of each protein. After excluding the terminal residues, the respective correlations remained similar at 0.70, 0.69 and 0.83, compared with 0.71, 0.70 and 0.84 for the full domains (Fig. 2B). This indicates that the agreement between short- and long-timescale normalised RMSF was not primarily driven by terminal flexibility.

To further assess the reliability of short MD simulations, we compared RMSF calculated from experimental NMR structural ensembles with RMSF obtained from 1-ns MD simulations. We selected 239 NMR structures from the RCSB PDB, each containing at least 30 deposited models and 100 residues. NMR-based RMSF was calculated from all deposited models, whereas MD-based RMSF was obtained from 1-ns GROMACS simulations initiated from the first model of each NMR ensemble using the AMBER force field. Both RMSF profiles were normalised within each protein. The resulting profiles showed a positive correlation (Pearson *r* = 0.74; Fig. 2C).

To further evaluate sensitivity to initial structures, simulation engines, and force fields, a representative subset of 300 proteins was randomly sampled from the test set. This subset comprised 67,973 residues, providing sufficient data for further comparisons while maintaining computational tractability for re-simulation. To verify that this subset accurately reflects the broader dataset, the timescale comparison was repeated exclusively for these 300 structures. The correlation between 1-ns and 100-ns RMSF profiles for this specific subset remained high (Pearson *r* = 0.87).

The sensitivity of RMSF profiles to the choice of initial structural model was next examined. Normalised RMSF profiles computed from AlphaFold2-derived domain structures were compared with those obtained from the corresponding crystallographic domains. The two sets of normalised RMSF values exhibited a strong correlation (Pearson *r* = 0.81). Robustness with respect to MD engines and force-field choice was further assessed using the representative subset. Normalised RMSF profiles derived from simulations performed with OpenMM (Version 7.0) and GROMACS (Version 2023.3) [1, 9] using identical AMBER parameters for 1 ns were highly correlated (Pearson *r* = 0.78). Similarly, comparisons between different force fields in GROMACS (AMBER versus OPLS) yielded strong agreement (Pearson *r* = 0.79) (Fig. 2D).

### 3.3 Prediction Accuracy and Benchmark Evaluation of ProtFlap

The predictive performance of ProtFlap was evaluated using the nonredundant dataset of 7,878 CATH domains. During cross-validation across ten independent random splits, the model demonstrated stable predictive accuracy (Supplementary Table S3). Evaluation on the validation sets yielded a mean root-mean-square error (RMSE) of 0.74 and Pearson correlation coefficient of 0.65. Variance across the ten splits was marginal, indicating robust model convergence independent of specific data partitioning. Performance on the held-out 10% independent test set was consistent with the cross-validation metrics.

ProtFlap was subsequently benchmarked against four established predictors: RMSF-Net [36], ProDy (v2.6.1) [48], BioEmu [25], and inverse weighted contact number (iWCN) [14, 27, 47]. For this comparative evaluation, the final ProtFlap model was trained on the complete training and validation pool and evaluated on the independent test set. To assess variation across proteins, Pearson correlations and RMSE values were calculated separately for each protein (Fig. 3(A–B)). In the continuous prediction of normalised RMSF, ProtFlap achieved the highest mean Pearson correlation coefficient 0.66 *±* 0.19) and the lowest mean RMSE (0.73 *±* 0.16) among all evaluated methods.

**Figure 3:**
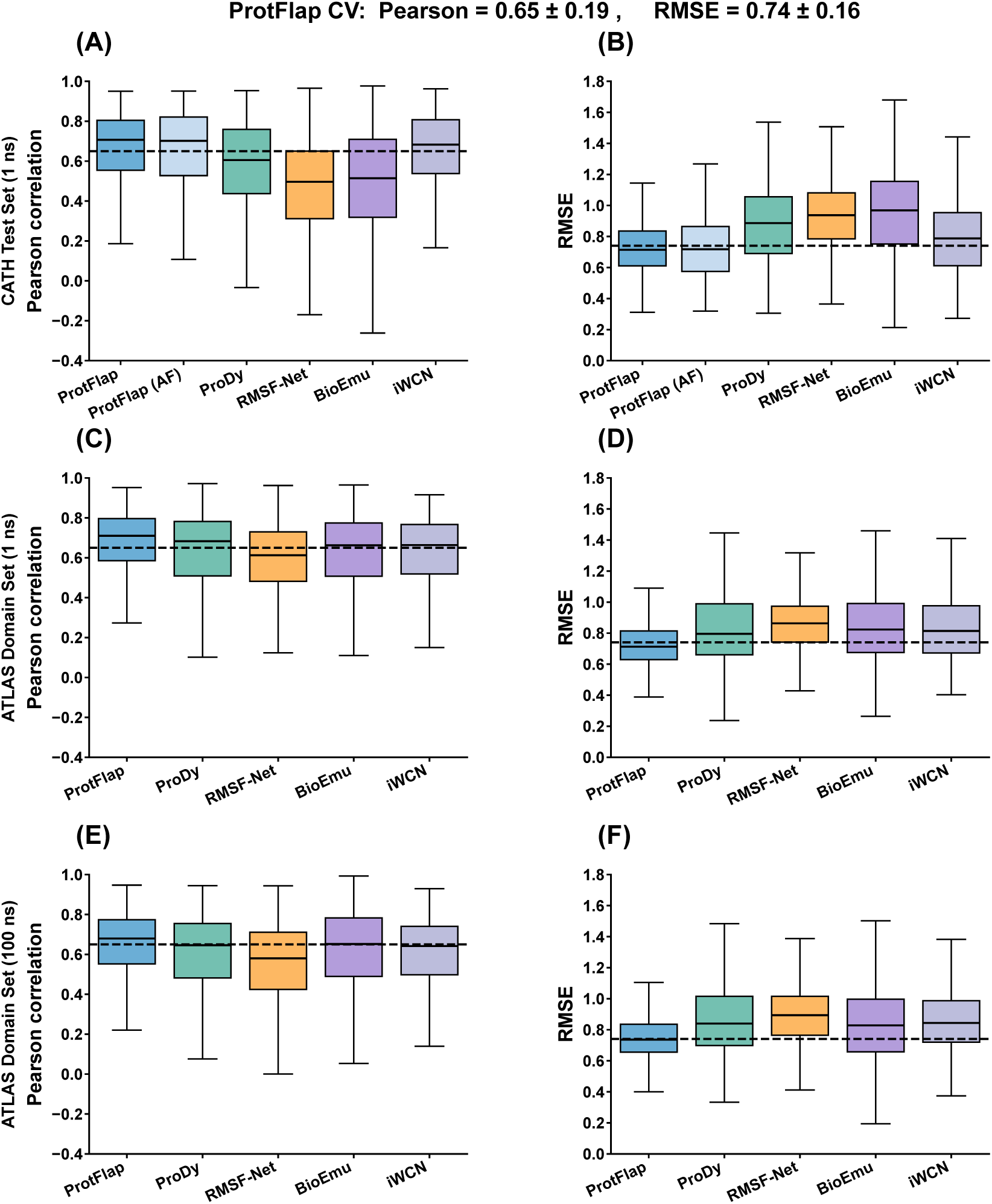
Predictive performance and benchmarking of ProtFlap. (A–B) Distributions of per-domain (A) Pearson correlation coefficients and (B) root-mean-square errors (RMSE) on the independent CATH test set, comparing ProtFlap, ProtFlap using AlphaFold-derived structures (AF), ProDy, RMSF-Net, BioEmu, and iWCN. (C–F) Corresponding comparisons on the independent ATLAS domain set using RMSF derived from 1-ns (C,D) and 100-ns (E,F) MD simulations as reference. Boxes indicate the interquartile range, with central lines showing the median. Dashed horizontal lines indicate the mean cross-validation performance of ProtFlap.

To assess robustness to input structure origin, ProtFlap was additionally evaluated using AlphaFold-generated structural models (AF Model) for the test set. Performance on these AI-predicted structures remained nearly identical to the experimental inputs (Pearson *r* = 0.66 *±* 0.20, RMSE = 0.72 *±* 0.19). Interestingly, the simple iWCN descriptor achieved the second-highest mean Pearson correlation (0.65 *±* 0.20), closely approaching ProtFlap, although its RMSE was higher (0.79 *±* 0.23).

To further assess generalisability, we evaluated ProtFlap on the independent ATLAS dataset (domain wise) (Fig. 3(C–D)). ProtFlap maintained the highest mean Pearson correlation (0.66 *±* 0.19) and the lowest mean RMSE (0.73 *±* 0.15) among the evaluated methods. In comparison, iWCN achieved a mean Pearson correlation of 0.63 *±* 0.19 and an RMSE of 0.83 *±* 0.21. ProDy, BioEmu, and RMSF-Net also showed lower predictive accuracy than ProtFlap on this dataset. We also evaluated the methods against normalised RMSF derived from 100-ns ATLAS simulations (Fig. 3(E–F)). ProtFlap retained the highest mean Pearson correlation (0.64) and lowest mean RMSE (0.76) among the evaluated methods, indicating that its predictive performance remained consistent when longer MD simulations were used as the reference. The distributions of per-protein performance and the corresponding summary statistics are presented in Fig. 3 and Supplementary Table S4, respectively.

We analyzed per-protein performance across CATH classes and protein lengths and compared the best-performing and failure cases (top and bottom 10%). The failure group showed a higher proportion of All-*α* proteins, which may be more challenging to predict due to their characteristically high structural stability and relatively limited conformational variability (Supplementary Fig. S2).

### 3.4 Feature Importance, Ablation and Structural Analysis

Node-level feature importance analysis using SHAP [28] revealed that atomic energy terms, which capture the local thermodynamic constraints on a residue, dominated model predictions. The Van der Waals-related energy component (EV), a direct proxy for local packing density and steric clashes, contributed approximately 30% of the total importance (Fig. 4A). Other energetic features governing local geometry, including electrostatic (EC), torsional (ET), and bond (EB, ES) terms, provided consistent but more modest contributions. Empirical distributions for all six energetic features demonstrated significant separation between flexible and non-flexible residues, confirming that the model successfully leverages fundamental physical forces for classification (Cramér–von Mises test; Fig. 4B and Supplementary Fig. S3).

**Figure 4:**
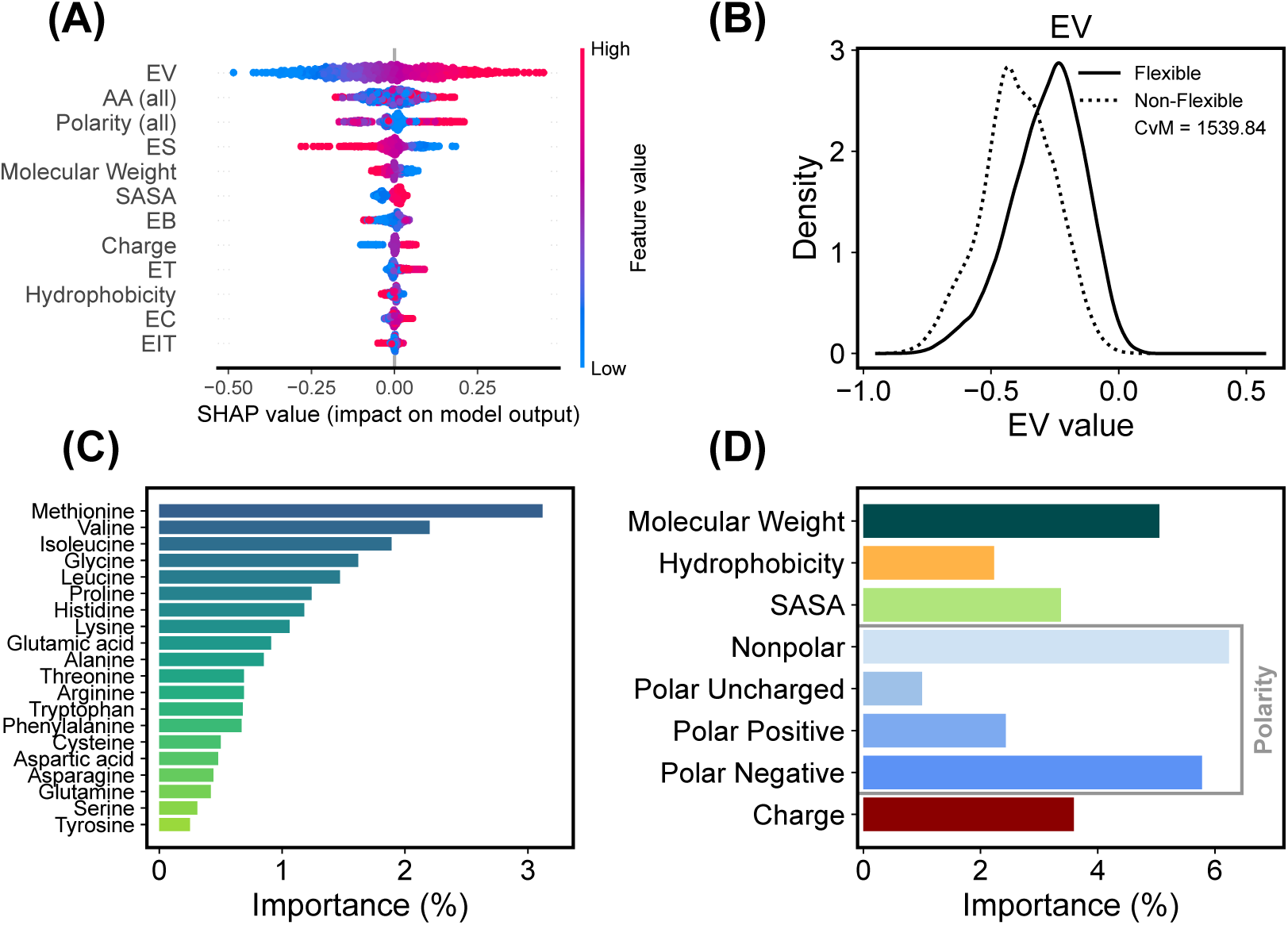
Feature importance analysis of ProtFlap predictions. (A) SHAP (Shapley Additive Explanations) summary plot detailing the relative contribution of node-level features to flexibility prediction. Colour gradients indicate raw feature values (red: high, blue: low). (B) Empirical distribution of the Van der Waals-related energy component (EV) for flexible versus non-flexible residues, annotated with the Cramér–von Mises test statistic. (C) SHAP-based feature importance rankings for individual amino acid identities. (D) Relative importance contributions of sequence-derived physicochemical and structural features, including molecular weight, hydrophobicity, solvent-accessible surface area (SASA), polarity, and net charge.

Amino acid identity formed the second most influential feature group. Residues that classically dictate structural core stability (hydrophobic valine and methionine) and those governing solvent-exposed interactions (charged arginine and aspartic acid) were major contributors to the classification (Fig. 4C). Solvent-accessible surface area (SASA) and polarity descriptors further stratified these predictions, physically reflecting the distinct local environments of buried cores versus exposed surfaces (Fig. 4D).

The sensitivity of ProtFlap to the energy-calculation protocol was further examined by comparing two configurations: AMBER99SB with GB-HPMF and OPLS with ACE (Supplementary Fig. S4). Both configurations showed comparable predictive performance (Pearson *r* = 0.66 and 0.63; RMSE = 0.73 and 0.75, respectively), with slightly better results for AMBER99SB/GB-HPMF. SHAP analysis showed that van der Waals energy (EV) remained among the most influential features in both configurations, although the relative importance of other features varied. These findings suggest that the contribution of energetic information is maintained across the two protocols examined.

To further assess the contribution of different feature groups, we also performed ablation studies (Supplementary Table S5). Removing the atomic-energy features reduced the mean Pearson correlation from 0.65 for the full model to 0.58 for the Sequence–Structure model, indicating that energetic information contributes to RMSF prediction beyond conventional sequence and structural descriptors. The Energy-Only model achieved a correlation of 0.59, suggesting that atomic-energy features alone provide useful predictive information but are less effective than their combination with sequence and structural features. The No-Solvation model showed intermediate performance (*r* = 0.62), indicating an additional contribution from solvation-related information. Replacing the GCN with an MLP reduced the correlation to 0.53, suggesting that graph-based information propagation also contributes to prediction accuracy. The full model achieved the lowest RMSE (0.74) among the evaluated configurations.

### 3.5 Structural Characteristics and Prediction Performance Across Protein Groups

Analysis of secondary structure assignments derived from DSSP [20] showed that loop regions, which lack regular stabilising constraints, exhibited the highest flexibility, *α*-helices demonstrated intermediate flexibility, and *β*-strands, typically constrained by extensive inter-strand hydrogen bond networks, were predominantly rigid (Fig. 5A). Model performance across different protein fold architectures was assessed using CATH classifications [23]. Predictive accuracy was largely consistent across mainly *α*, mainly *β*, and *α*/*β* architectures (Fig. 5B). Predictions for mainly *β* folds yielded the highest predictive performance, recording an AUROC of 0.81 and a PRAUC of 0.74. Architectures classified as *α*+*β* recorded the lowest predictive accuracy among the evaluated structural groups(Fig. 5(C–D)).

**Figure 5:**
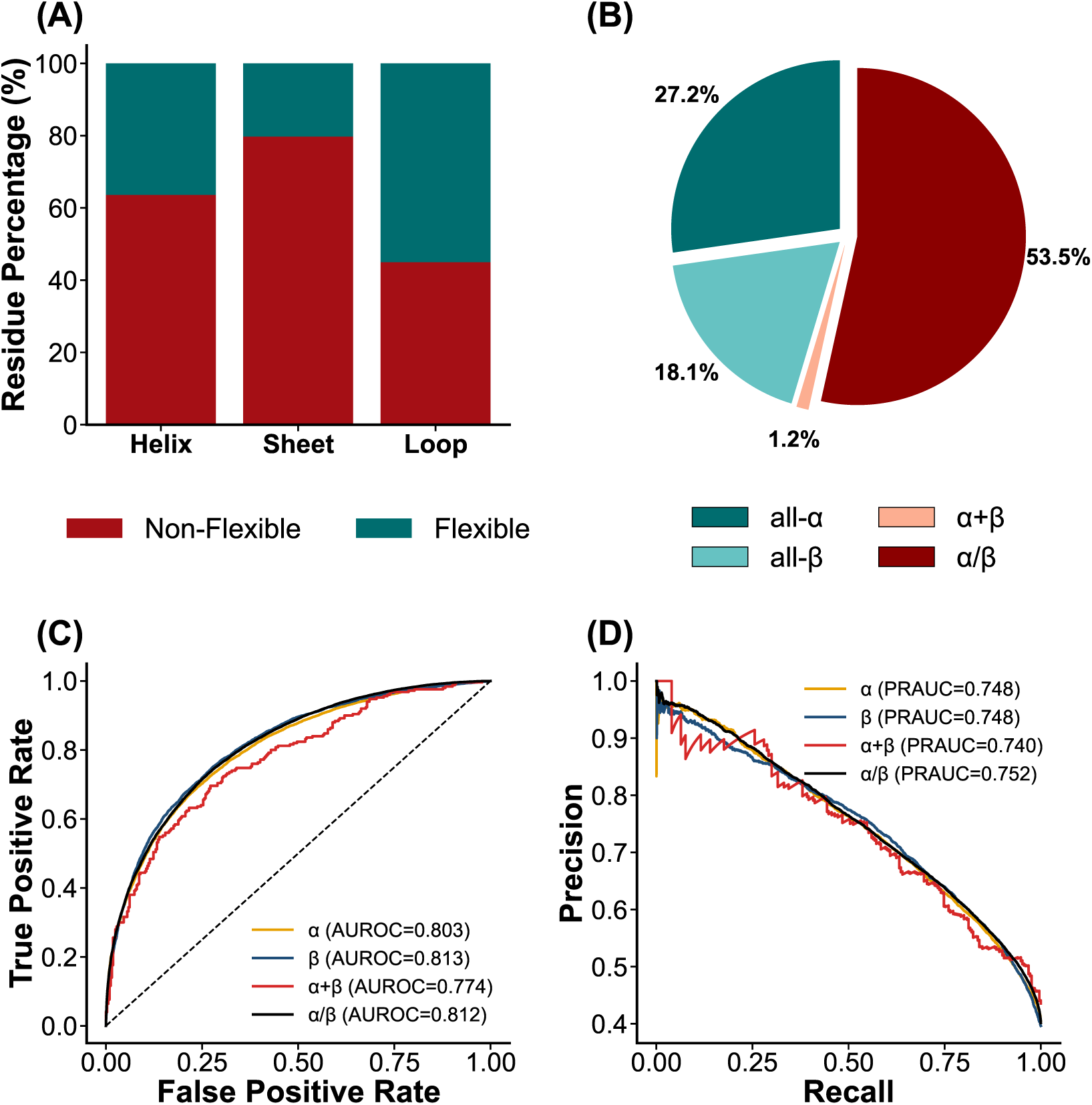
ProtFlap predictions and performance stratified by structural classifications. (A) Percentage distribution of flexible versus non-flexible residues across DSSP-assigned secondary structure elements (helices, sheets, and loops). (B) Proportion of CATH structural architectures (all-*α*, all-*β*, *α*+*β*, and *α*/*β*) represented within the independent test set. (C–D) Receiver operating characteristic (ROC) curves (C) and precision-recall (PR) curves (D) evaluating binary classification performance across the four CATH structural classes.

To further examine variation in prediction accuracy, we analysed per-domain Pearson correlations and RMSE across structural classes and domain lengths (Supplementary Fig. S2). Performance was broadly similar across the major structural classes, whereas the *α* + *β* group showed a lower mean correlation (*r* = 0.57) and higher RMSE (0.81). Prediction performance also varied across domain lengths, with significant differences in Pearson correlation (*p* = 0.036) and RMSE (*p* = 0.018; Kruskal–Wallis test). Domains containing 400 or more residues generally showed lower correlations and higher errors. We further compared the best- and worst-performing 10% of domains, revealing differences in their length distributions and structural class composition (Supplementary Fig. S2E,F).

We also examined prediction accuracy in proteins containing disorder-associated regions (Supplementary Fig. S5). Using 28 PDB chains comprising 6,841 residues, ProtFlap achieved a Pearson correlation of 0.79 and an RMSE of 0.61. Within the 1,292 annotated disordered residues, ProtFlap maintained a high correlation (*r* = 0.82), although the RMSE increased to 0.79. ProtFlap performed better than ProDy, BioEmu, RMSF-Net and iWCN in both whole-protein and disordered-region comparisons. The higher RMSE within disordered regions suggests that these regions remain more difficult to predict quantitatively.

## 4 Discussion

We present ProtFlap, a structure-based deep learning framework that estimates residue-level protein flexibility without requiring explicit dynamical simulation. Through the integration of atomic energy features with structural descriptors, ProtFlap links the local thermodynamic environment of a static protein structure to its intrinsic mobility.

A central finding of this study is that the local atomic energy network, extracted from a single energy-minimised snapshot, contains strong predictive signals for residue flexibility. Feature attribution analysis identified van der Waals-related energy (EV) as the most influential feature in the SHAP analysis. This aligns with biophysical principles: regions with loose local packing and fewer steric constraints experience weaker Van der Waals interactions, thereby permitting greater thermal motion. The strong distributional separation of energetic features between flexible and non-flexible residues confirms that ProtFlap exploits these physical constraints rather than relying on structural memorisation.

The model successfully captures the complex interplay between amino acid identity, solvent exposure, and secondary structure. Hydrophobic residues are correctly identified as predominantly rigid, reflecting their role in dense core packing. Conversely, charged and polar residues contribute to flexibility primarily through surface exposure. Furthermore, ProtFlap accurately reproduces macroscopic biophysical trends, identifying loops as highly flexible and *β*-strands as inherently rigid. The high predictive accuracy observed for mainly *β* architectures is a direct consequence of this; *β*-sheets are stabilised by extensive, regular hydrogen-bond networks that severely restrict local fluctuations [39].

While ProtFlap demonstrates robust generalisation across most fold classes, predictive accuracy was lower for *α*+*β* architectures.This may partly reflect the unequal representation of fold classes within the CATH-derived training dataset. Machine learning models are inherently constrained by their training distributions, and the relative scarcity of complex *α*+*β* topologies likely restricted the model’s ability to map their specific energetic environments. Future iterations of the framework will require stratified sampling of structural databases to ensure balanced representation across all major fold classifications.

A critical advantage of the ProtFlap framework is its robustness to varied input conditions. The target metric, normalised RMSF, proved highly consistent across different simulation lengths, molecular dynamics engines, and force fields. More importantly, ProtFlap achieved comparable accuracy when applied to AlphaFold-derived domain models and experimental crystal structures. This indicates that the energetic environments within predicted structures are sufficiently self-consistent to estimate dynamics.

By reframing protein flexibility as an emergent property of the static energetic landscape, ProtFlap bridges the gap between structural biology and molecular dynamics. It provides a physically grounded, interpretable method for flexibility prediction, with an average processing time of 52.3 s per protein structure, including energy minimisation, feature extraction and inference (Supplementary Table S6).

## Acknowledgments

We sincerely thank The Brain Korea 21 (BK21) FOUR Program funded by the Ministry of Education (MOE) and National Research Foundation (NRF) of Korea (No. 2023-0827-03).

## Author contributions statement

H.L. and Y.C. conceived the experiment(s), H.L. conducted the experiment(s), H.L. and Y.C. analysed the results. H.L. and Y.C. wrote and Y.C. reviewed the manuscript.

## Data availability

The software is available on GitHub (https://github.com/yoonjoolab/ProtFlap). The test set data and predictions are provided on (https://github.com/yoonjoolab/ProtFlap/tree/main/test_set).

