## Supplementary for "ProtFlap: Structure-based Deep Learning Prediction of Protein Flexibility from Atomic Energy Networks"

### ProtFlap: Predicting Residue-Level Protein Flexibility from Atomic-Energy-Informed Interaction Graphs

**Supplementary Fig. S1. Optimisation of the Cα distance cutoff.** Five Cα distance cutoffs (5, 10, 15, 20 and 30 Å) were evaluated using 10-fold cross-validation. Bars show mean Pearson correlation coefficients, with error bars indicating standard deviations. The 15 Å cutoff gave the highest mean correlation (r = 0.65) and was selected for the final model.

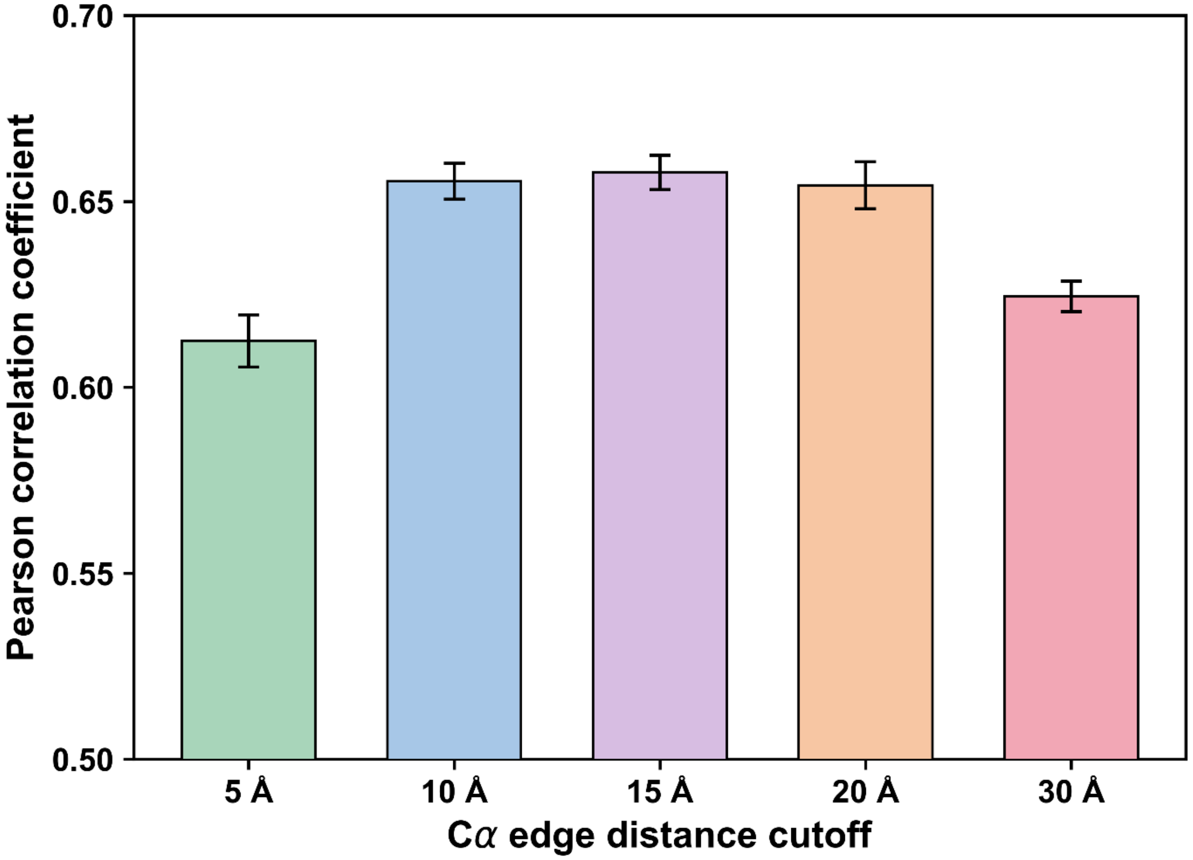

**Supplementary Figure S2. ProtFlap prediction performance across CATH fold classes and protein length groups.** (A) Mean Pearson correlation across CATH fold classes: All-α (0.64), All-β (0.67), α/β (0.67), and α+β (0.57). (B) Mean RMSE across CATH fold classes: All-α (0.75), All-β (0.73), α/β (0.73), and α+β (0.81). (C) Pearson correlation across five protein length groups, with an overall per-protein mean of 0.66 and median of 0.70. (D) RMSE across five protein length groups, with an overall per-protein mean of 0.73 and median of 0.72. Prediction performance varied significantly across protein length groups for Pearson correlation (Kruskal–Wallis, $p = 0.036$) and RMSE ($p = 0.018$). (E) Cumulative length distributions of the best-performing and failure groups (top and failure 10%). (F) CATH composition of the best-performing and failure groups: All-α (26.92% vs. 32.05%), All-β (19.23% vs. 17.95%), α/β (53.85% vs. 48.72%), and α+β (0% vs. 1.28%).

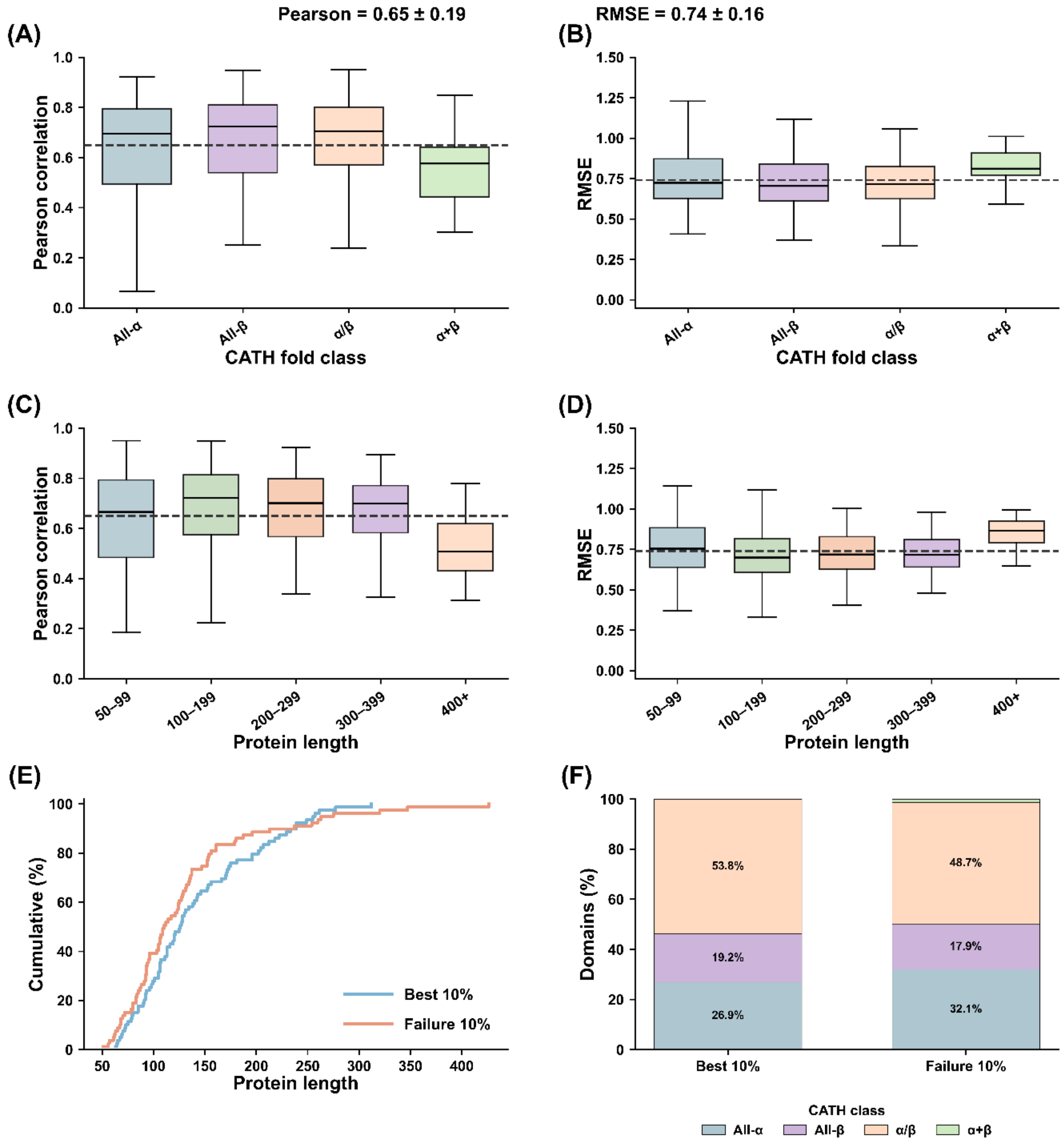

**Supplementary Fig. S3. Distributions of atomic energy terms for flexible and non-flexible residues.** Cramér–von Mises (CvM) statistics were calculated for the six most influential atomic energy terms to assess their discriminative power between flexible and non-flexible residues. The van der Waals-related energy component (EV) exhibited the highest CvM statistic. The remaining energy terms (EB, ET, EIT, EC, and ES) provided consistent, albeit more modest, contributions to flexibility discrimination. Notably, the solvation energy term (ES) reflects residue–solvent interactions modelled using the GB-HPMF implicit solvation scheme during energy minimisation.

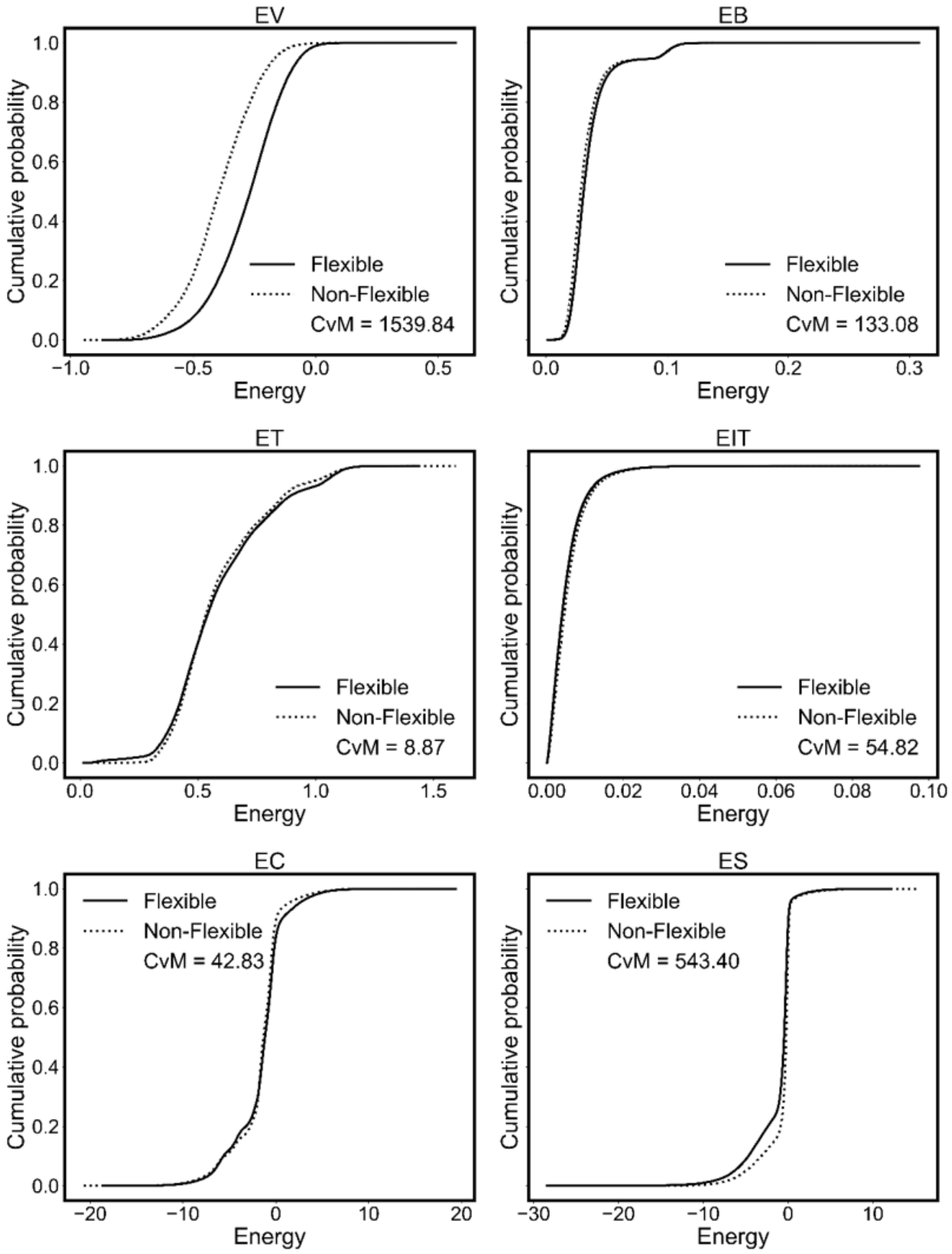

**Supplementary Fig. S4. Comparison of force-field configurations and their effects on prediction performance and feature importance**. ProtFlap was trained using two TINKER/SOLVATE force-field configurations: ACE (OPLS) and GB-HPMF (AMBER99SB), using the same training and test datasets. **(A)** Comparison of prediction performance on the held-out test set based on Pearson correlation, root mean square error (RMSE), GB-HPMF showed slightly better performance, with a higher mean Pearson correlation (0.66 vs. 0.63) and lower RMSE (0.73 vs. 0.75) than ACE. **(B)** Comparison of SHAP-based feature importance across top features for the two force-field configurations, illustrating differences in the relative contributions of individual features to model predictions.

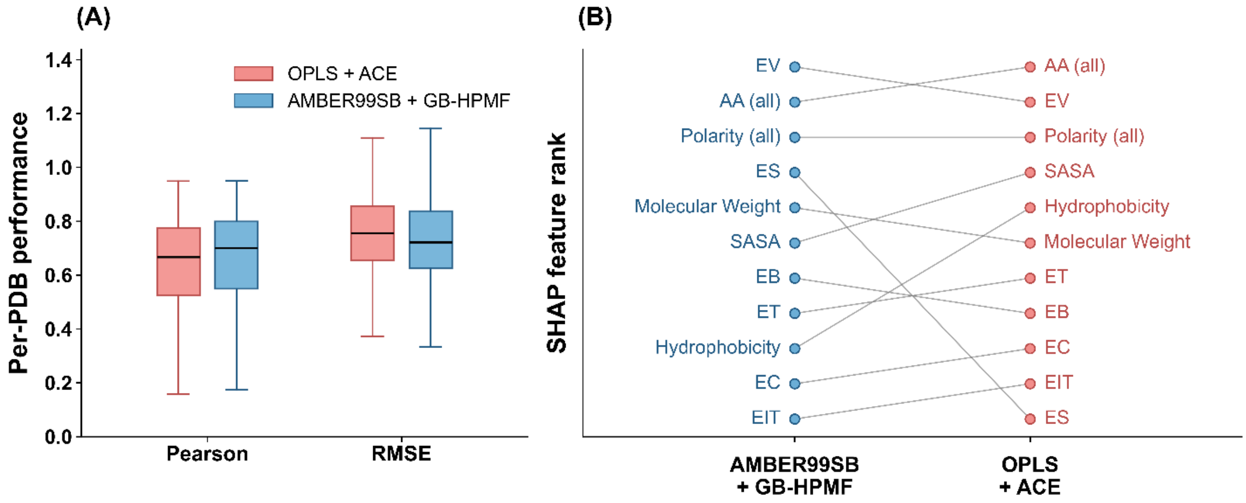

**Supplementary Figure S5. Comparison of protein flexibility prediction methods across whole proteins and disordered regions.** Pooled residue-level Pearson correlation coefficients and root mean square errors (RMSE) were calculated for ProtFlap, ProDy, BioEmu, RMSF-Net, and iWCN using 28 matched PDB chains. The analysis included 6,841 residues across the whole proteins and 1,292 residues within their corresponding disordered regions. ProtFlap achieved the highest Pearson correlation for both whole-protein (r = 0.79) and disordered-region predictions (r = 0.82), with RMSE values of 0.61 and 0.79, respectively.

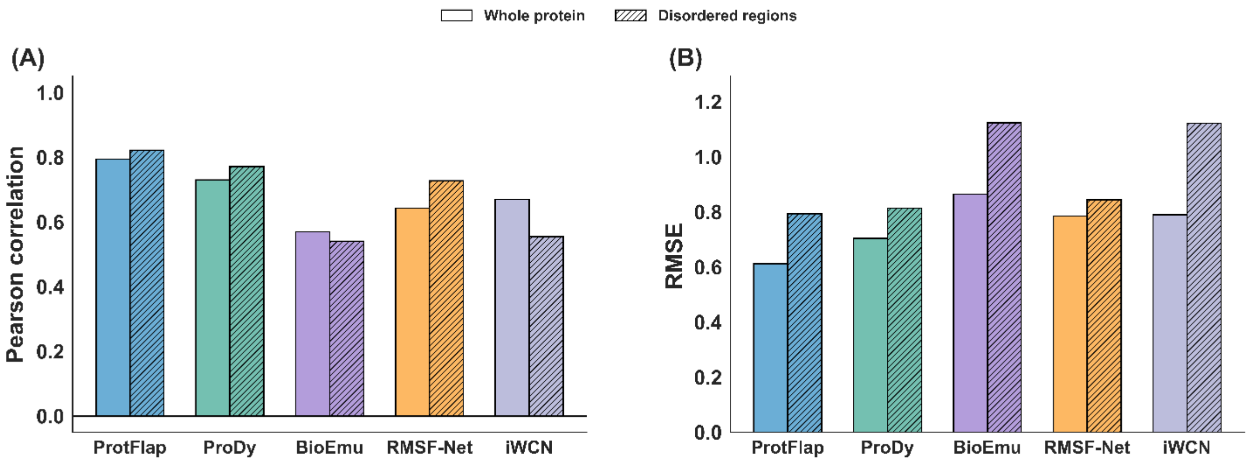

**Supplementary Table S1. Hyperparameter optimisation across evaluated graph neural network architectures.**

| Parameter | GCN | GIN | GraphSAGE |
|---|---|---|---|
| Best Validation Loss | 0.25 | 0.26 | 0.27 |
| Learning Rate | 0.0001 | 0.001 | 0.001 |
| Dropout | 0.16 | 0.20 | 0.05 |
| Weight Decay | 0.001 | 0.0001 | 0.0001 |
| Batch Normalization | True | True | True |
| Residual Connections | True | True | False |
| Activation Function | ELU | ReLU | ELU |
| Use Bias | False | False | True |
| Hidden Dimension | 64 | 64 | 256 |
| Number of Layers | 6 | 6 | 4 |

**Supplementary Table S2. Functional categorization of atomic energy terms computed using Tinker.**

| Category | Tinker Notation | Description |
|---|---|---|
| Bonded terms | EB | Bond stretch potential energy |
| | EA | Angle bend potential energy |
| | ET | Torsional potential energy |
| | EID | Improper dihedral potential energy |
| | EIT | Improper torsion potential energy |
| | EOPD | Out-of-plane distance potential energy |
| | EOPB | Out-of-plane bend potential energy |
| | EUB | Urey–Bradley potential energy |
| | EAT | Angle–torsion potential energy |
| | EBA | Stretch–bend potential energy |
| | EBT | Stretch–torsion potential energy |
| | ETT | Torsion–torsion potential energy |
| | EAA | Angle–angle potential energy |
| | EPT | π-system torsional potential energy |
| Nonbonded / environmental terms | EC | Charge–charge potential energy |
| | ECD | Charge–dipole potential energy |
| | ED | Dipole–dipole potential energy |
| | EM | Atomic multipole potential energy |
| | EP | Polarization potential energy |
| | ER | Pauli repulsion potential energy |
| | EV | van der Waals potential energy |
| | EDSP | Dampled dispersion potential energy |
| | ECT | Charge-transfer potential energy |
| | ERXF | Reaction-field potential energy |
| | ES | Solvation potential energy (GB, PB, or surface-area models) |
| Specialized / extra terms | ELF | Metal ligand-field potential energy |
| | EG | Geometric restraint potential energy |
| | EX | Extra term potential energy |

**Supplementary Table S3. Validation set per protein performance across ten independent random splits.**

| Fold | Pearson (Mean ± SD) | RMSE (Mean ± SD) |
|---|---|---|
| 1 | 0.65 ± 0.19 | 0.74 ± 0.15 |
| 2 | 0.64 ± 0.20 | 0.75 ± 0.16 |
| 3 | 0.65 ± 0.19 | 0.75 ± 0.15 |
| 4 | 0.65 ± 0.19 | 0.74 ± 0.15 |
| 5 | 0.65 ± 0.20 | 0.74 ± 0.16 |
| 6 | 0.65 ± 0.19 | 0.74 ± 0.15 |
| 7 | 0.65 ± 0.19 | 0.74 ± 0.16 |
| 8 | 0.65 ± 0.19 | 0.74 ± 0.16 |
| 9 | 0.65 ± 0.19 | 0.74 ± 0.16 |
| 10 | 0.65 ± 0.19 | 0.74 ± 0.16 |
| **Overall** | **0.65 ± 0.19** | **0.74 ± 0.16** |

**Supplementary Table S4. Comparative evaluation of ProtFlap and existing methods for protein flexibility prediction on the CATH test set and ATLAS dataset**

| Dataset | Metric | ProtFlap | ProtFlap (AF) | ProDy | RMSF-Net | BioEmu | iWCN |
|---|---|---|---|---|---|---|---|
| **CATH Test Set (1 ns)** | Pearson | 0.66 ± 0.19 | 0.66 ± 0.20 | 0.58 ± 0.22 | 0.47 ± 0.25 | 0.49 ± 0.27 | 0.65 ± 0.20 |
| | RMSE | 0.73 ± 0.16 | 0.72 ± 0.19 | 0.88 ± 0.24 | 0.93 ± 0.22 | 0.95 ± 0.28 | 0.79 ± 0.23 |
| **ATLAS (1 ns)** | Pearson | 0.66 ± 0.19 | — | 0.63 ± 0.21 | 0.59 ± 0.18 | 0.62 ± 0.21 | 0.63 ± 0.19 |
| | RMSE | 0.73 ± 0.15 | — | 0.83 ± 0.24 | 0.87 ± 0.19 | 0.84 ± 0.23 | 0.83 ± 0.21 |
| **ATLAS (100 ns)** | Pearson | 0.64 ± 0.20 | — | 0.60 ± 0.22 | 0.55 ± 0.21 | 0.61 ± 0.22 | 0.60 ± 0.20 |
| | RMSE | 0.76 ± 0.15 | — | 0.86 ± 0.23 | 0.90 ± 0.20 | 0.84 ± 0.24 | 0.86 ± 0.21 |

**Supplementary Table S5: Feature and architectural configurations evaluated in ProtFlap ablation analysis.**

| Model | Sequence | Structure | Energy | Solvation | GCN | MLP | Pearson | RMSE | No. of Features |
|---|---|---|---|---|---|---|---|---|---|
| **Energy-Only** | | | ✓ | | ✓ | ✓ | 0.59 | 0.80 | 28 |
| **Sequence–Structure** | ✓ | ✓ | | | ✓ | ✓ | 0.58 | 0.81 | 31 |
| **No Solvation** | ✓ | ✓ | ✓ | | ✓ | ✓ | 0.62 | 0.78 | 59 |
| **Full Feature MLP** | ✓ | ✓ | ✓ | ✓ | | ✓ | 0.53 | 0.84 | 59 |
| **Full Model** | ✓ | ✓ | ✓ | ✓ | ✓ | ✓ | 0.65 | 0.74 | 59 |

**Supplementary Table S6. Comparison of average computational runtime per protein structure across ProtFlap and benchmark methods.**

| Method | Average runtime per PDB |
|---|---|
| ProtFlap | 52.3 sec |
| ProDy (GNM) | 0.04 sec |
| RMSF-Net | 8.28 sec |
| BioEmu | 15 min 2 sec |
| iWCN | 0.022 sec |
